# Mechanism of duplex unwinding by coronavirus nsp13 helicases

**DOI:** 10.1101/2020.08.02.233510

**Authors:** Xiao Hu, Wei Hao, Bo Qin, Zhiqi Tian, Ziheng Li, Pengjiao Hou, Rong Zhao, Sheng Cui, Jiajie Diao

**Affiliations:** NHC Key Laboratory of Systems Biology of Pathogens, Institute of Pathogen Biology, Chinese Academy of Medical Sciences and Peking Union Medical College, Beijing 100730, China; Department of Cancer Biology, University of Cincinnati, Cincinnati, OH 45267, USA

**Keywords:** MERS-CoV, SARS-CoV-2, nsp13 helicase, smFRET, unwinding mechanism

## Abstract

The current COVID-19 pandemic urges in-depth investigation into proteins encoded with coronavirus (CoV), especially conserved CoV replicases. The nsp13 of highly pathogenic MERS-CoV, SARS-CoV-2, and SARS-CoV exhibit the most conserved CoV replicases. Using single-molecule FRET, we observed that MERS-CoV nsp13 unwound DNA in discrete steps of approximately 9 bp when ATP was used. If another NTP was used, then the steps were only 4 to 5 bp. In dwell time analysis, we detected 3 or 4 hidden steps in each unwinding process, which indicated the hydrolysis of 3 or 4 dTTP. Based on crystallographic and biochemical studies of CoV nsp13 helicases, we modeled an unwinding mechanism similar to the spring-loaded mechanism of HCV NS3 helicase, although our model proposes that flexible 1B and stalk domains, by allowing a lag greater than 4 bp during unwinding, cause the accumulated tension on the nsp13-DNA complex. The hinge region between two RecA-like domains in SARS-CoV-2 nsp13 is intrinsically more flexible than in MERS-CoV nsp13 due to the difference of a single amino acid, which causes the former to induce significantly greater NTP hydrolysis. Our findings thus establish a blueprint for determining the unwinding mechanism of a unique helicase family.

1. When dTTP was used as the energy source, 4 hidden steps in each individual unwinding step after 3 - 4 NTP hydrolysis were observed.
2. An unwinding model of MERS-CoV-nsp13 which is similar to the spring-loaded mechanism of HCV NS3 helicase, except the accumulation of tension on nsp13/DNA complex is caused by the flexible 1B and stalk domains that allow a lag of 4-bp in unwinding.
3. Comparing to MERS-CoV nsp13, the hinge region between two RecA-like domains in SARS-CoV-2 nsp13 is intrinsically more flexible due to a single amino acid difference, which contributes to the significantly higher NTP hydrolysis by SARS-CoV-2 nsp13.

## Introduction

Coronaviruses (CoVs) pose unprecedented threats to human health and the global economy, as powerfully demonstrated by the current pandemic caused by coronavirus disease 2019 (COVID-19). The third major CoV outbreak in the past two decades, COVID-19 has triggered devastating effects dwarfing those of severe acute respiratory syndrome (SARS) in 2002 and 2003 (1,2) and Middle East respiratory syndrome (MERS) in 2012 (3).

SARS-CoV-2 (4), the causative agent of COVID-19, is the seventh human CoV (5), one with approximately 80% nucleotide sequence similarity with SARS-CoV. Although the shared sequence variation is more pronounced in the structural (i.e., envelope E, membrane M, nucleocapsid N, and spike S) and accessory proteins (i.e., ORF3a or 3b, 6, 7a or 7b, 8, and 10) of SARS-CoV-2 than SARS-CoV, its nonstructural proteins are highly conserved (6). In fact, the central components of the viral RNA replication transcription complex’s (RTC) nonstructural proteins (nsp)—that is, nsp13 (i.e., helicase) and nsp12 (i.e., RNA-dependent RNA polymerase)—are more than 98% identical. Perhaps more remarkable, the nsp13 helicase encoded by three highly pathogenic CoVs—SARS-CoV, MERS-CoV, and SARS-CoV-2—shares 84% to 99% sequence similarity, thereby making it a potential wide-spectrum antiviral target. As a member of the SF1B helicase family, nsp13 unwinds RNA or DNA duplexes with 5’-to-3’ directionality and hydrolyzes both NTP and dNTP (7,8); it also hosts 5’-triphosphatase activity. In previous work, by determining the crystal structure of MERS-CoV nsp13 (PDB ID: 5WWP), we revealed that nsp13 generally mirrors the domain organization of nidovirus helicases, whereas the individual domains (i.e., CH, RecA1–1B and RecA2) of nsp13 reflect the organization of cellular Upf1-like helicases (9, 10).

Mechanisms of CoV nsp13 have mostly been investigated by using ensemble methods. SARS-CoV sp13 prefers a substrate with long 5’ single-stranded (ssDNA) tails or gaps for duplex unwinding, in which case multiple nsp13 molecules loading on the 5’ ssDNA regions can enhance the processivity of DNA unwinding (11). Following the results of a hydrogen–deuterium exchange assay, a translocation mechanism of nsp13 was proposed (12), one that details the grasp and release of ssDNA relay between the RecA1–1B and RecA2 domains of nsp13, which correlate with three transition states of ATP hydrolysis. That model largely reflects the paradigmatic mechanism of the SF1B helicase RecD2 (13). Another intriguing observation was that the unwinding reaction catalyzed by nsp13 contained lags, and such lagging became more evident as DNA duplex became longer. Using the rapid chemical quench flow method, Sarafianos and colleagues detected intermediates during dsDNA unwinding by SARS-CoV nsp13 and estimated an average kinetic step size of 9.3 bp per step (14). However, the mechanism underlying the lagged unwinding was not elaborated.

Because the structural and biochemical characterizations of MERS-CoV and SARS-CoV nsp13 are available, and because SARS-CoV-2 nsp13 differs from SARS-CoV nsp13 by only one conservative substitution (i.e., I570V), we investigated the kinetic mechanism that allows two nsp13 helicases to unwind DNA. Our findings not only elucidate that unique helicase’s mechanism but also may contribute to developing antiviral agents to combat COVID-19, its potential resurgence, and the outbreak of similar viral diseases in the future.

## Results

### MERS-CoV Helicase Rapidly Unwound DNA

In our previous biochemical characterization, the way in which MERS-CoV nsp13 unwound an 18-bp DNA duplex containing a 5’ overhang depended upon NTP hydrolysis. Similarly, SARS-CoV nsp13 was able to unwind a 15-bp partial DNA duplex with the same 5’ overhang. To elucidate their unwinding mechanism, we designed single-molecule fluorescence resonance energy transfer (smFRET) experiments (**Fig. 1A**). Using a biotin–NeutrAvidin interaction, we immobilized partial double-stranded DNAs (dsDNAs) with 5’ ssDNA tails to the imaging surface with biotin at the 5’ termini of the ssDNA overhangs. The partial dsDNA consisted of an 18-base top strand, labeled at the 3’ end by Cy5 as the acceptor, and a 37-base bottom strand, labeled at the ssDNA–dsDNA junction by Cy3 as the donor. After pre-incubating 20 nM of full-length MERS-CoV nsp13 protein with the DNA substrate, we injected 4 mM of ATP into the system to initiate the reaction. The corresponding FRET values represented different unwinding states. Before we added ATP, nsp13 bound to the ssDNA portion of the partial duplex, most likely at the ssDNA–dsDNA junction; thus, the sustained proximity of the donor and acceptor generated high FRET values. Following the addition of ATP, nsp13 began to translocate on the bottom strand toward the 3’ end and unzipped the DNA duplex. Once the donor and the acceptor had separated, FRET values gradually decreased. We visualized the unwinding process by total internal reflection fluorescent (TIRF) microscopy; representative images of donor and acceptor channels within 1 min after ATP injection appear in **Fig. 1B**.

**Figure 1.**
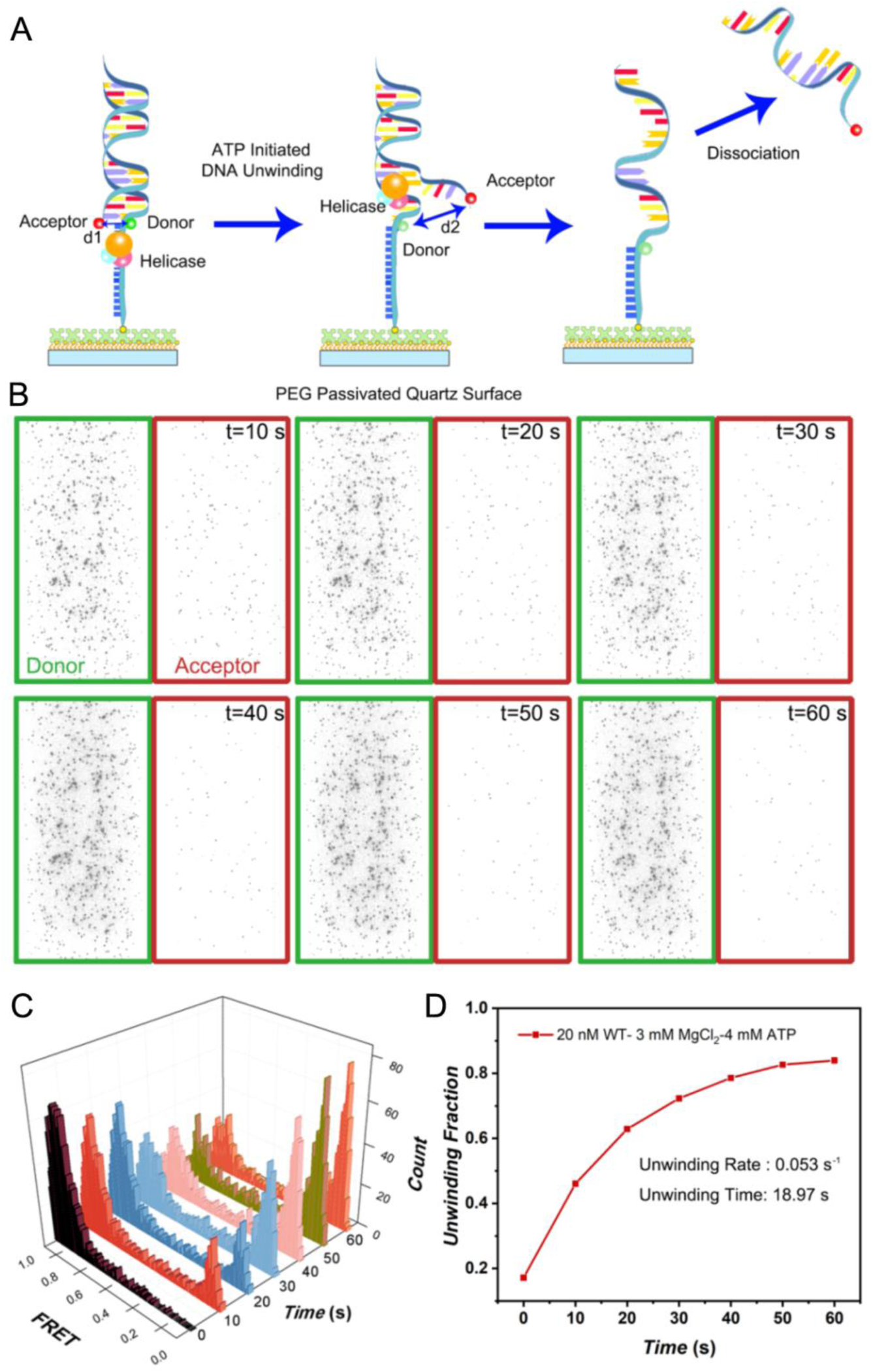
MERS-CoV nsp13 helicase unwound most double-stranded DNAD (dsDNA) in 1 min. (a) Schematic illustration of DNA unwinding by MERS-CoV nsp13 helicase using adenosine triphosphate (ATP). (b) Fluorescent images of DNA in corresponding donor (Cy3) and acceptor (Cy5) channels 1 min after the addition of ATP. (c) Histograms of FRET efficiency as a function of time showing the population of unwinding DNA (i.e., low FRET) and no unwinding DNA (high FRET). (d) Unwound DNA (i.e., high FRET population) as a function of time measured from the histograms in (c); an overall exponential fit to the curve demonstrated an unwinding rate, *R*, of 0.053 s^-1^ and an unwinding time of 18.97 s.

We evaluated the corresponding FRET values of the immobilized dsDNAs at different time points after injecting them with ATP (**Fig. 1C)**. All histograms revealed a high FRET peak followed by a low one, which respectively corresponded to the original dsDNAs and unwound DNAs. At time 0 (i.e., before ATP was added), most of the dsDNAs exhibited high FRET values with negligible low FRET peaks. Following the injection of ATPs, the high FRET peaks decreased with time, whereas the low FRET peaks increased. Next, we plotted the unwound population according to time (**Fig. 1D**). At the initial phase of the reaction, MERS-CoV nsp13 unwound rather rapidly but decelerated with time, most likely due to the system’s consumption of DNA substrate and ATP. Approximately 45% of the total dsDNA population unwound in the first 10 s, and approximately 80% had unwound after 1 min. The overall unwinding rate, *R*, and unwinding time, *T*, were determined by fitting the evolution of the unwinding population with a single exponential function, for values of *R* = 0.053 s^-1^ and *T* = 18.97 s at the selected experimental condition.

### MERS-CoV nsp13 Exhibited Single and Multiple Steps in Unwinding

Under our initial experimental conditions, MERS-CoV nsp13 unwound the DNA duplex in a single step in most cases, although multistep unwinding also occurred. **Fig. 2A** shows representative traces of different unwinding processes. In traces to the left, both processes showed a single FRET decrease indicating single-step unwinding; in traces to the right, two FRET jumps indicated two working steps with 9 bp per step. The stable intermediate FRET values suggest that MERS-CoV nsp13 paused the unwinding of the DNA duplex under certain conditions, which aligns with a finding from a previous ensemble study on SARS-CoV nsp13-catalyzed unwinding reactions (14). Therein, Sarafianos and colleagues discovered the presence of intermediates in the unwinding of a DNA duplex by SARS-CoV nsp13, and using the pre-steady state kinetic assay, they estimated that the unwinding step was approximately 9.3 bp, which is remarkably similar to unwinding step size of MERS-CoV nsp13 in our study.

**Figure 2:**
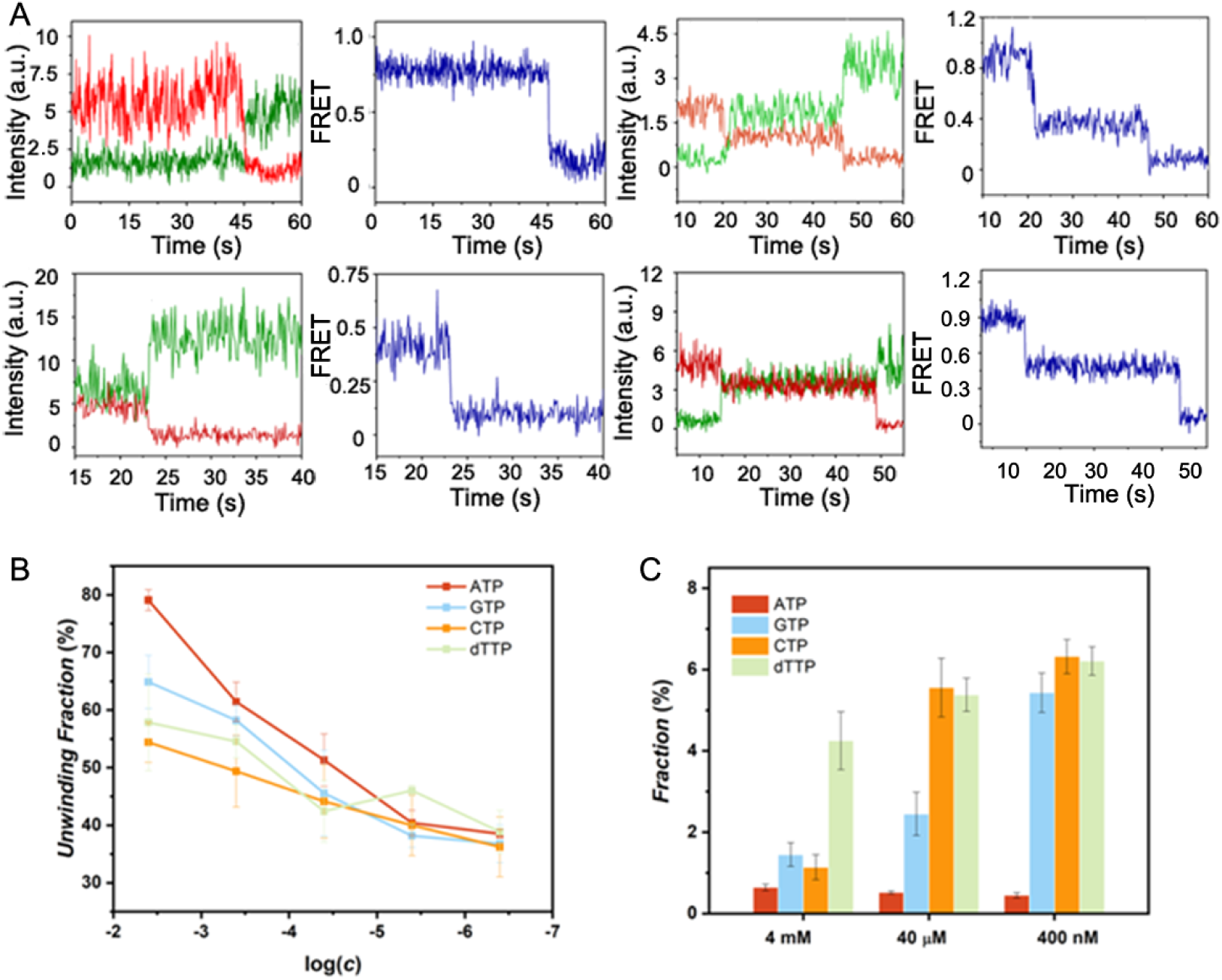
The MERS-CoV nsp13 helicase unwound dsDNA in single or multiple steps, although more multistep processes appeared with the decreasing of unwinding rate. (a) Representative fluorescent intensity and corresponding FRET efficiency time traces of single-step (left) and multistep (right) unwinding processes. (b) Unwinding rates of MERS-CoV nsp13 decreased with the concentrations of chemical energy (i.e., ATP, GTP, CTP, and dTTP). (c) More multistep unwinding processes appeared at lower concentrations of chemical energy. Incidence of multistep unwinding is plotted as the function of NTP concentrations; results were summarized from at least 1,000 traces.

To resolve multistep unwinding, we decelerated the unwinding by supplying different chemical energy sources at low concentrations. Helicases have different preferences in relation to NTPs; for example, Hepatitis C virus (HCV) NS3 hydrolyses all eight canonical NTPs in the order of preference (d)ATP > (d)CTP > UTP or dTTP > (d)GTP (15). Therefore, we used various NTPs (i.e., ATP, GTP, CTP, and dTTP) to fuel the unwinding reactions and indeed achieved far slower unwinding rates by using those NTPs as energy sources or at lower concentrations (i.e., 4 mM to 400 nM), as shown in **Fig. 2B**. The unwound population decreased from approximately 80% to 35% when we reduced the ATP concentrations from 4 mM to 400 nM. Other energy sources decreased unwinding rates at lower concentrations. Although MERS-CoV nsp13 clearly preferred ATP over the other NTPs at high concentrations (i.e., 40 µM to 4 mM), that preference became subtler at lower concentrations (i.e., 400 nM to 4 µM). As expected, decreasing the unwinding rate increased the incidence of the multistep unwinding event. When the concentration of GTP, CTP, or dTTP decreased to 400 nM, approximately 6% of the unwinding events displayed a multistep unwinding process (**Fig. 2C**). However, that result never occurred when we used ATP as the energy source, not even when its concentration was 400 nM and the unwinding rate was as low as that of CTP, GTP, and dTTP.

### MERS-CoV nsp13 Unwound the dsDNA in 4 Discrete Steps

The transition density plot (**Fig. 3A**) was summarized from 140 transitions with FRET changes of approximately 0.25. Four peaks with enter/exit FRET emerged at approximately 0.97/0.75, 0.76/0.51, 0.50/0.26, and 0.25/0.07. The transition density plot confirmed that the 0.25 FRET length was the smallest unwinding step observed in experiments, and 4 discrete unwinding steps emerged in the unwinding process of 18 bp with 400 μM of dTTP.

**Figure 3.**
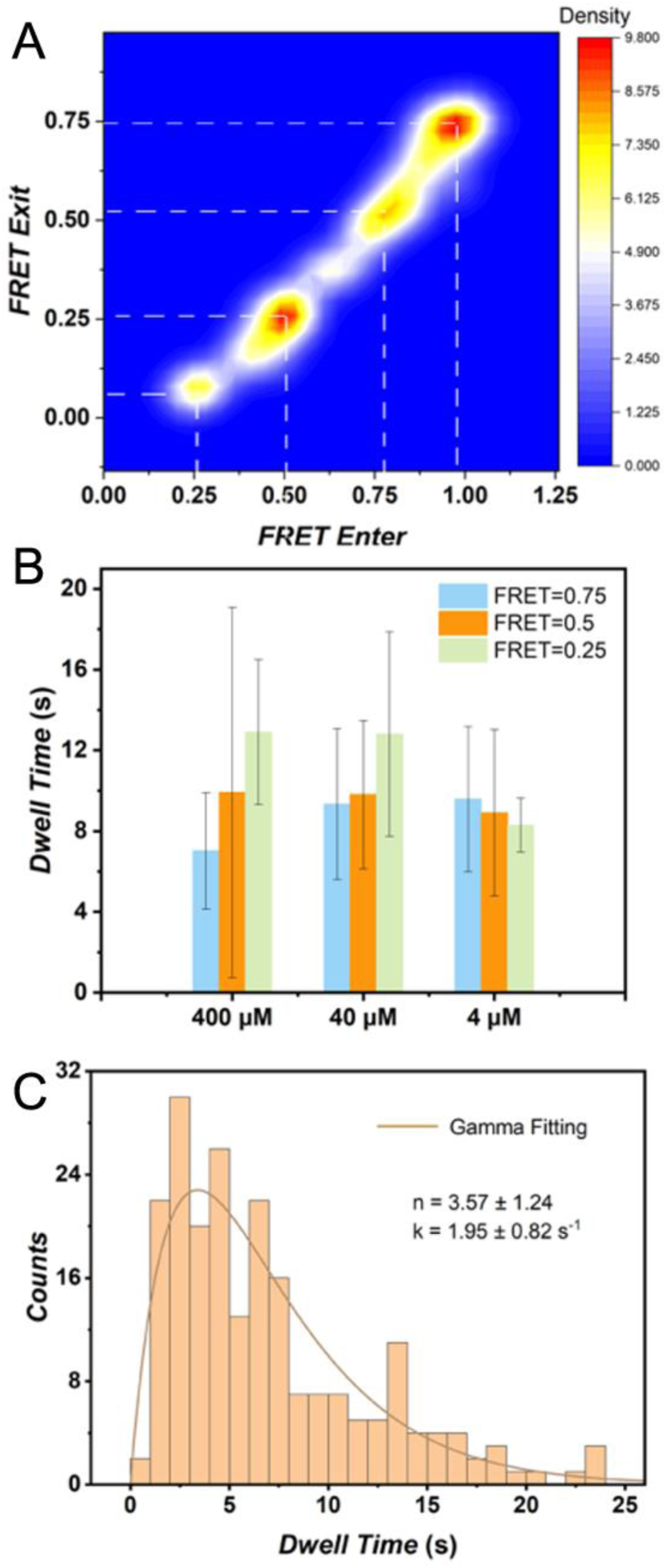
The MERS-CoV nsp13 helicase unwound the 18-bp dsDNA in 4 discrete steps. (a) FRET values obtained from 140 traces were combined to make the transition density plot. (b) Dwell time of each status at different concentrations of chemical energy; 216 molecules with a multistep unwinding process were used to calculate dwell time. (c) Gamma distribution fitting of the collected dwell times at each plateau’s pause duration.

The trace curves revealed that MERS-CoV nsp13 unwound the dsDNA in stepwise fashion. Specifically, MERS-CoV nsp13 lingered at a specific unwinding step (i.e., FRET state) before commencing the next one. The term *dwell time* (i.e. FRET exit time minus FRET enter time) is used to describe how long MERS-CoV nsp13 stayed at a specific FRET state. Dwell time at each FRET state was evaluated from 216 molecules with a multistep unwinding process, and the results (**Fig. 3B**) indicate that MERS-CoV nsp13 lingered at every FRET state for approximately 6.94 ± 5.06 s before commencing the next step. However, dwell time did not show obvious changes with lower concentrations of energy sources.

The dwell time histogram of individual steps appears in **Fig. 3C**. Because the dwell time histogram did not follow a single-exponential decay, irreversible Poissonian steps were hidden in the observed individual steps. Gamma distribution, *t*^*n*–1^ *exp*(–*kt*), was used to fit the dwell time histogram, and the calculated *n* and *k* values that emerged were 3.57 ± 1.24 and 1.95 ± 0.82 s^-1^, respectively, indicating 3 or 4 hidden irreversible Poissonian steps, presumably due to the hydrolysis of single dTTP, with an identical rate *k* of approximately 1.95 s^-1^. Based on those results, our unwinding model for MERS-CoV helicase maintains that, after 3 or 4 successive dTTP hydrolysis events, an abrupt 4-bp separation occurred.

### Single Point Mutations Affected the Unwinding Rate of MERS nsp13

To gain further insights into the unwinding mechanism, we selected 13 residues, all invariant in SARS-CoV and SARS-CoV-2 nsp13, from across the nucleic acids binding groove of MERS-CoV nsp13 for a mutagenesis study (**Fig. 4**). We predicted that the selected residues would contact ssDNA based on the modeled structures of MERS-CoV nsp13-ssDNA (9) and SARS-CoV-2 nsp13-ssDNA. The mutations included S310D, H311D, T359D, N361D, A362D, and P408D on the RecA1 domain; R178D on the 1B domain; and Y515D, N516D, T532D, D534A S535D, and R560D on the RecA2 domain (**Fig. 4A & 4B**). Special features of the selected residues were twofold. First, all selected residues had structural counterparts in EVA nsp10, whose involvement in DNA binding has been confirmed by X-ray crystallography, as shown in **Table 1**. Second, the selected residues were located on conserved DNA binding motifs; S310 and H311 from motif Ia and residue P408 from motif III formed a conserved pocket similar to the motif-Ia-pocket in the 1A domain of RecD2, which generally may accommodate a DNA base. Residues T359, N361, and A362 from the loop between β18 and β19 were located at the entrance of the ssDNA binding channel, near the 3’ end of the modeled ssDNA. The position of that loop was equivalent to the pin device in RecD2 helicase, albeit far shorter in nsp13 and without forming a rigid β-hairpin structure. The part from Y515 to N516, from the loop between β23 and α14, was located at the 5’ end terminus of the modeled ssDNA. T532, D534, and S535 from motif V formed a short helix below the phosphodiester backbone of the modeled ssDNA. Meanwhile, R560 from motif VI was one of few residues that may interact with the base and sugar moiety of the ssDNA. R178 from the 1B domain interacted with the modeled ssDNA, which played a role in fixing ssDNA to the 1B, thus causing a lag in unwinding, as according to our postulation (**Fig. 5**). The conserved threonine pair (i.e., with one threonine on each RecA-like domain) appeared in the structurally characterized SF1A helicases (i.e., Rep, UvrD and PcrA), SF1B helicase (RecD2), and SF2 helicase; HCV NS3 is also available in MERS-CoV and SARS-CoV nsp13 helicases, T359 and T532. The threonine pair served as the reference point for understand the translocation mechanism, such that when ATP was bound to a helicase, the threonine residues were 2 nt apart, and when ATP was absent, they were 3 nt apart (16). That finding supports the widely accepted theory of 1-nt translocation coupling to 1 ATP hydrolysis. In our nsp13-ssDNA model representing the ATP-absent situation, the threonine pair is shown to be 3 nt apart, which also supports the theory. In that light, determining the structures of nsp13 DNA and RNA in the presence and absence of NTP or its analogues would contribute critical validation to the theory.

**Table 1.**
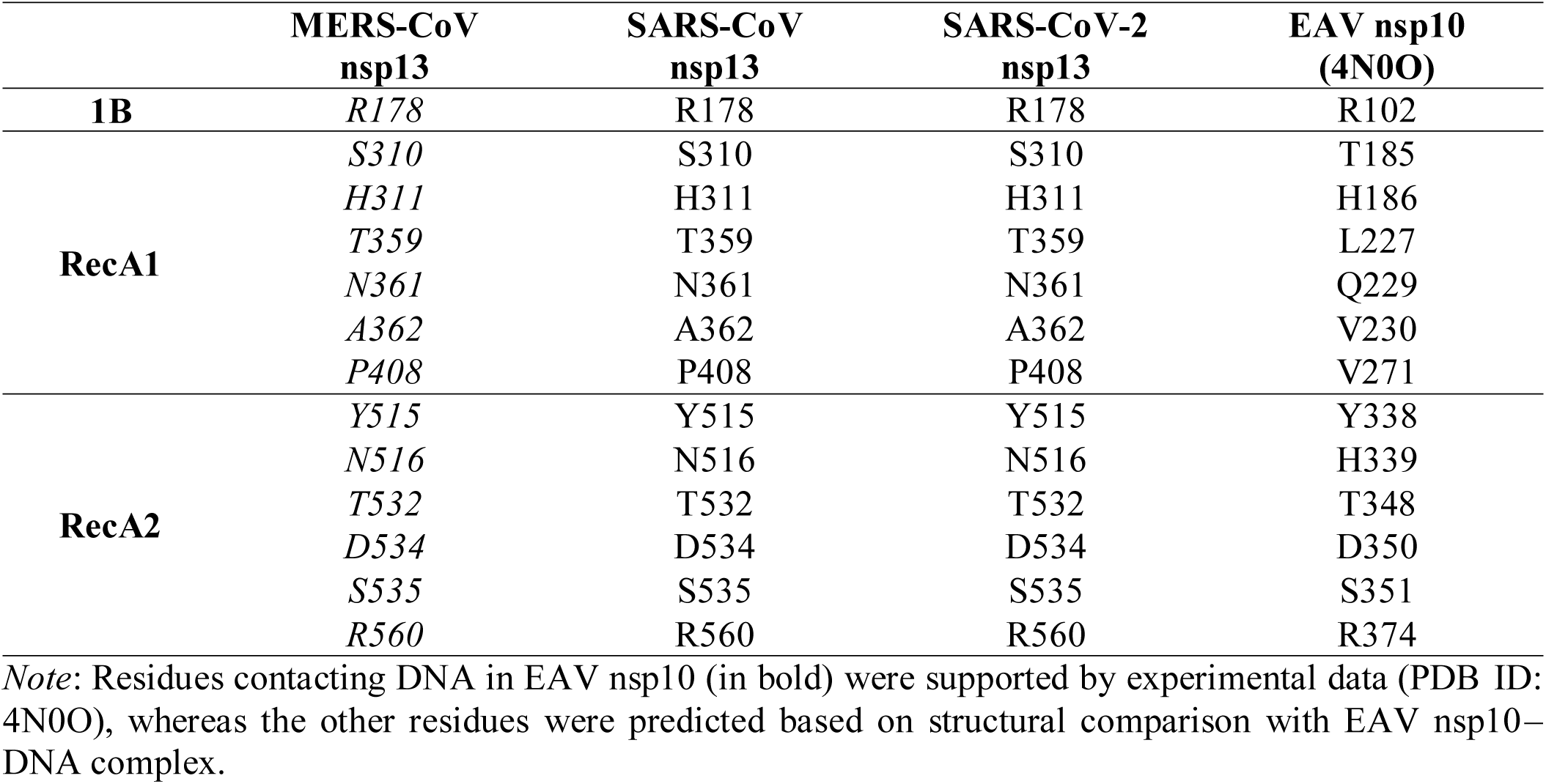
Residues predicted to contact DNA

|  | **MERS-CoV**<br>**nsp13** | **SARS-CoV**<br>**nsp13** | **SARS-CoV-2**<br>**nsp13** | **EAV nsp10**<br>**(4N0O)** |
| --- | --- | --- | --- | --- |
| **1B** | *R178* | R178 | R178 | R102 |
| **RecA1** | *S310* | S310 | S310 | T185 |
|  | *H311* | H311 | H311 | H186 |
|  | *T359* | T359 | T359 | L227 |
|  | *N361* | N361 | N361 | Q229 |
|  | *A362* | A362 | A362 | V230 |
|  | *P408* | P408 | P408 | V271 |
| **RecA2** | *Y515* | Y515 | Y515 | Y338 |
|  | *N516* | N516 | N516 | H339 |
|  | *T532* | T532 | T532 | T348 |
|  | *D534* | D534 | D534 | D350 |
|  | *S535* | S535 | S535 | S351 |
|  | *R560* | R560 | R560 | R374 |

*Note:* Residues contacting DNA in EAV nsp10 (in bold) were supported by experimental data (PDB ID: 4N0O), whereas the other residues were predicted based on structural comparison with EAV nsp10–DNA complex.

**Figure 4.**
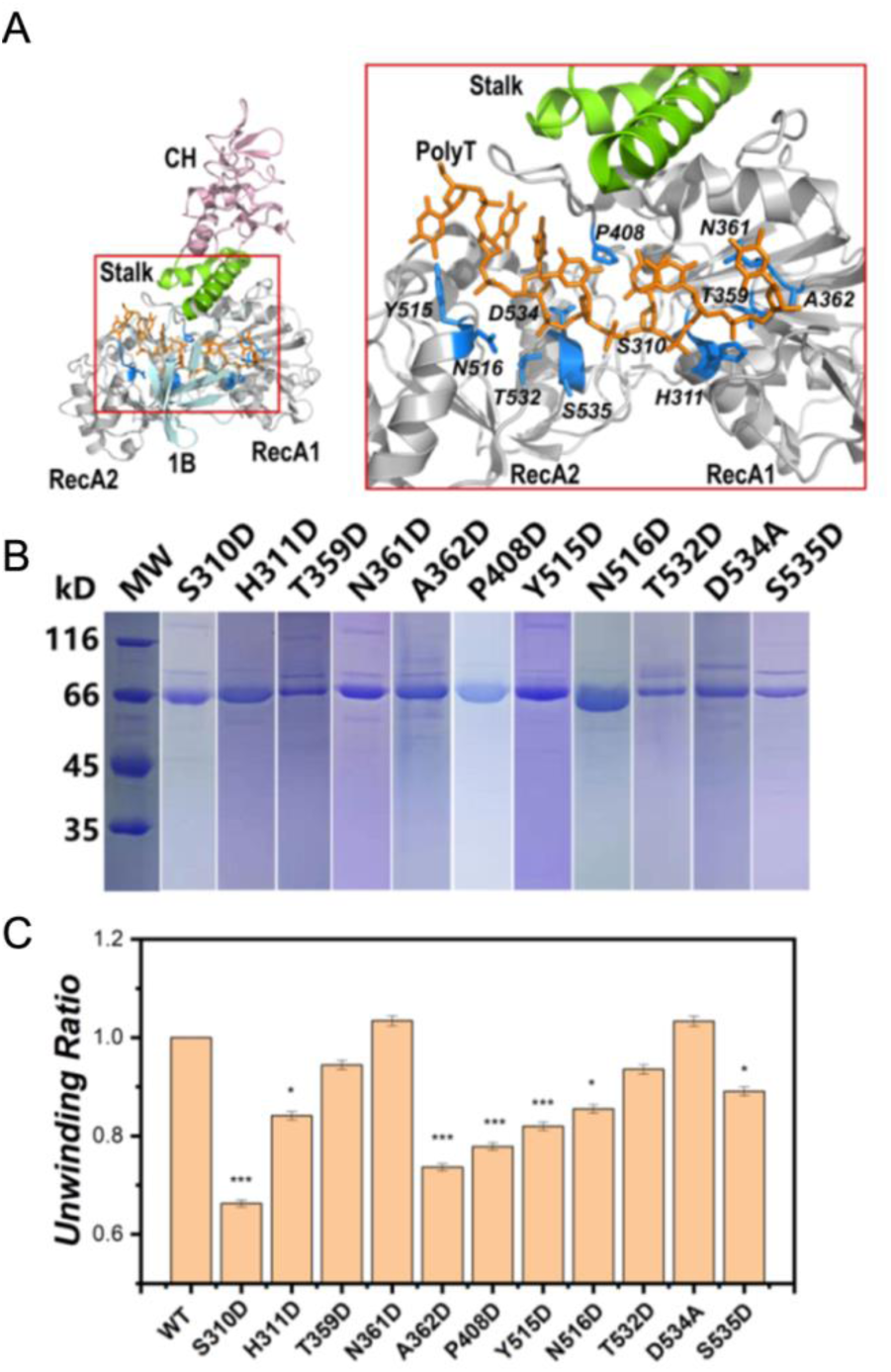
Single point mutations in the RecA1 and RecA2 domains affected the unwinding activity of MERS-CoV nsp13 helicase. (a) Structure-guided mutagenesis. Left: Ribbon model of MERS-CoV nsp13 with a modeled poly(dT) ssDNA strand. Domains of nsp13 are colored differently: CH domain in pink, stalk domain in green, 1B domain in cyan, and helicase core (i.e., RecA1 and RecA2) in gray. The poly(dT), in the orange stick model, was modeled by superimposing MERS-CoV nsp13 with EAV nsp10-ssDNA complex (PDB ID: 4N0O). To the right, a magnified view of the boxed area on the left depicts the putative RNA binding groove of MERS-CoV nsp13. Residues predicted to contact DNA appear in the blue stick model and are labeled in italics. Substitutions were introduced to each residue, which yielded 11 helicase mutants: S310D, H311D, T359D, N361D, A362D, P408D, Y515D, N516D, T532D, D534A, and S535D. The unwinding activities of the mutants were evaluated with sm FRET in (c). (b) SDS-PAGE gel image of 11 MERS-CoV nsp13 mutants. (c) Normalized unwinding rates of the WT and the different single point mutants. Student’s *t* test was used to determine the statistical significance between the WT and the mutants. * *p* < .05, ** *p* < .01, *** *p* < .005.

**Figure 5.**
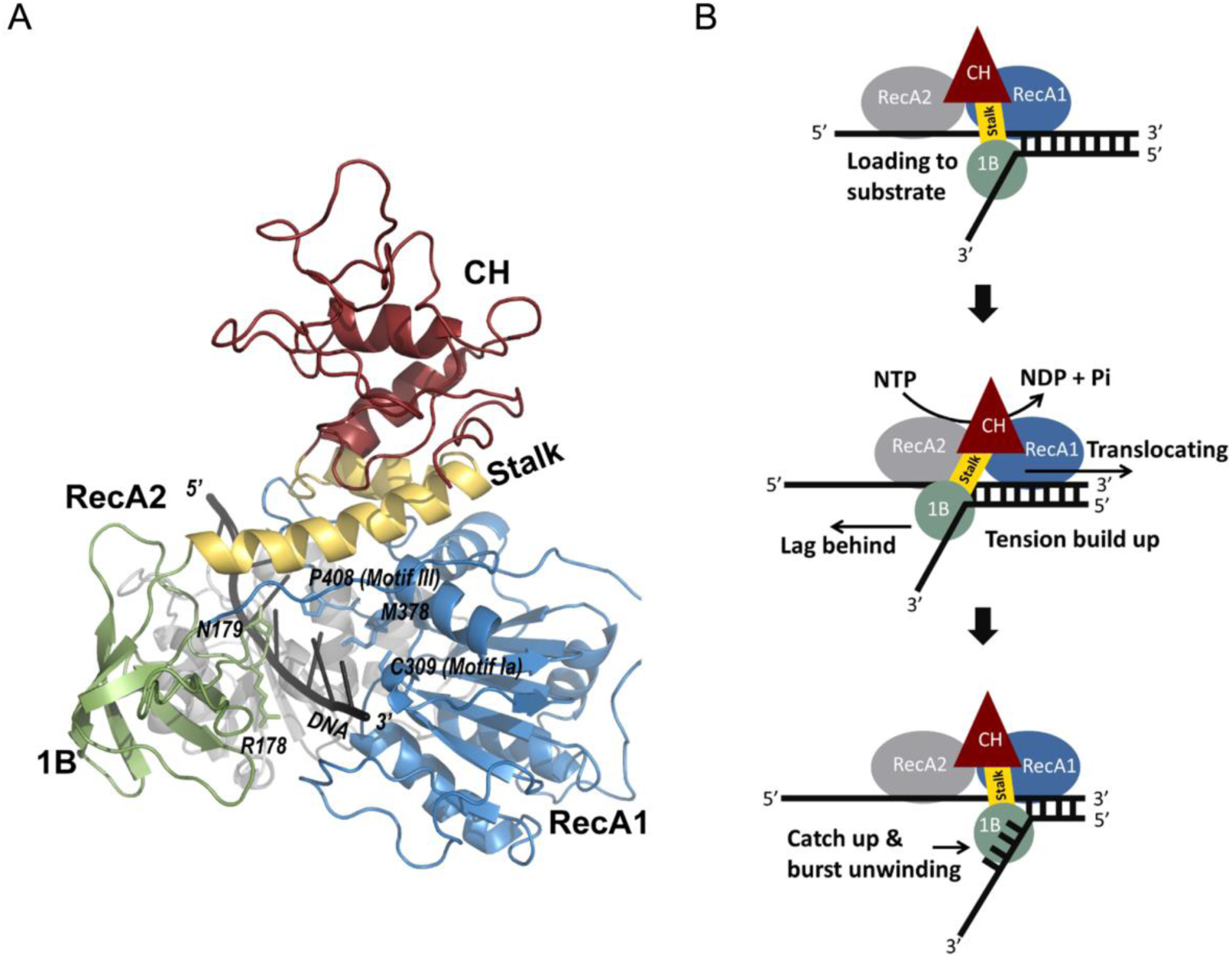
A model of CoV nsp13 unwinding. (A) A modeled structure of MERS-CoV nsp13 complexed with ssDNA. Individual domains are colored differently: CH domain in red, stalk domain in yellow, 1B domain in green, RecA1 domain in blue, and RecA2 domain in gray. DNA is in black. Key residues constituting the conserved domain between motif Ia and motif III appear in a stick model. Residues from 1B domain, which contacted the DNA on the opposite side, also appear in another stick model. (B) Model of CoV nsp13 translocating on DNA and the unwinding duplex with a 4-nt lag.

Using smFRET, we evaluated the unwinding rate of those mutants, except for R178D and R560D due to insolubility, and compared them to the wild-type (WT) level. Most mutations were detrimental to the unwinding activity (**Fig. 4C**). Compared with the WT nsp13, the loss in unwinding rate of S310D, A362D, P408D, and Y515D was extremely significant (*p* < .005), while the rates of H311D, N516D, and S535D were also significant (*p* < .01), whereas the rates of T359D, N361D, T532D, and D534A were not. The significant loss of activity caused by residue substitutions seems to reflect their vital role in unwinding. For example, residue P408 constituted one side of a pocket formed between motif Ia and III on the RecA1 domain. Designated “motif Ia pocket,” that pocket has been shown to play a vital role in DNA translocation, not only in SF1B helicase RecD2 (13) but also in the SF1A helicase, PcrA (17,18). During ssDNA translocation, a DNA base occupies the motif Ia pocket of RecD2, which acts as a physical block to grasp DNA while the 2A and 2B domains slip past. Because MERS-CoV nsp13 employs a similar mechanism, P408D mutation might have disrupted the hydrophobic interaction between DNA base and the motif Ia pocket, hence the impaired DNA translocation. Residue A362 is located on a loop between β18 and β19 at the entrance of the nucleic acid binding channel, where the ssDNA–dsDNA juncture is supposed to bind. Overlapping with the RecD2 structure, the equivalent region of the β18-to- β19 loop in RecD2 is a so-called “pin domain” functioning as a rigid device to split open the DNA–RNA duplex. Because A362 may engage in opening duplexes, its substitution by an aspartate might have hindered duplex separation instead of DNA translocation.

## Discussion

Available biochemical and structural evidence support a translocation model of CoV nsp13 in line with the canonical 5’-to-3’ translocation mechanism proposed for SF1B helicase (13). Nevertheless, prior to our study, the duplex-unwinding mechanism by nsp13 remained to be understood. Some helicases, including RecD2, RecG, and Hel308a (13,19,20), position a rigid pin or wedge device at the juncture of ssDNA–dsDNA to split open base pairs; however, CoV nsp13 lacks such device, or else the part exerting the analogous function within nsp13 has not been identified. Rao and colleagues reported a β19-to-β20 loop in SARS-CoV nsp13 equivalent to a β17-to-β18 loop in MERS-CoV nsp13, which hosts a handful of conserved, positively charged residues (e.g., Arg and Lys). They demonstrated that although that region is critical to nsp13-catalyzed unwinding activity, the loop only partially binds to the dsDNA substrate, not ssDNA alone. Revealing that region’s role in the unwinding process thus requires further study, especially to determine the structures of nsp13–dsDNA and nsp13–RNA complexes.

Our smFRET experiment clearly demonstrated that MERS-CoV nsp13-catalyzed dsDNA unwinding (i.e., an 18-bp partial duplex) has a discrete step size of 9 bp in the presence of ATP and from 4 to 5 bp when other NTP was used, which indicates the presence of a lag in unwinding. That result aligns exceedingly well with the findings of an ensemble study (14), in which researchers, using a Rapid Chemical Quench Flow instrument, estimated that SARS-CoV nsp13 unwinds dsDNA with discrete steps of 9.3 bp and a catalytic rate is 30 steps per second. The lag in duplex unwinding is reminiscent of the spring-loaded unwinding mechanism of the HCV NS3 helicase (HCV NS3-Hel) (21). HCV NS3 contains an aromatic residue W501 stacking against the 3’ terminus base of DNA, thereby fixing the DNA to the D3 domain. While two RecA-like D1 and D2 domains of the NS3-Hel translocate in the direction of 3’ to 5’ along DNA, the interaction between the D3 and DNA causes a lag in duplex unwinding. When the tension builds up to a breaking point, a sudden movement of D3 occurs with a concomitant burst of 3-bp unwinding.

We postulate that the 1B domain of CoV nsp13 may play a role similar to that of the D3 of NS3-Hel, albeit with the different mechanism. In our modeled structures of MERS-CoV nsp13 and SARS-CoV-2 nsp13 bound by ssDNA (9), the 1B domain and the RecA1 domain sandwich the 3’ portion of ssDNA. While the motif Ia pocket contacts the DNA on one side, the conserved residues 178R to 179N of the 1B domain contact the DNA on the other side. Fueled by the energy from NTP hydrolysis, the RecA1 and RecA2 translocate forward in a direction from 5’ to 3’, whereas the 1B remains fixed with the DNA. In contrast to the D3 of NS3-Hel, the 1B domain is distantly connected to the body of the helicase core via a rather long (i.e., approx. 20 Å) stalk domain. That unusually flexible architecture allows considerably greater interdomain rearrangement, thus the 1B–DNA interaction may survive the lagging at greater than 4 nt behind the translocating RecA1 and RecA2 domains.

Because SARS-CoV nsp13 and MERS nsp13 exhibit unmistakable similarity in structure (Cα rmsd =1.7 Å) and because SARS-CoV-2 nsp13 is nearly identical to SARS-CoV nsp13, the mechanism underlying the unwinding should be well preserved in all three nsp13 helicases. However, we discovered a notable difference at the hinge region (i.e., 438–445aa) between the RecA1 and RecA2 domains of nsp13. That hinge mediates their open-and-close rotation, which is essential to NTP hydrolysis and ssDNA translocation (12); thus, the flexibility of the hinge governs the activity of nsp13. MERS-CoV nsp13 contains residue S439 on the hinge, which is replaced by G439 in both SARS-CoV-2 nsp13 and SARS-CoV nsp13. Because glycine allows a wider range of dihedral angles than all non-glycine residues, the rotation between the RecA1 and RecA2 domains of SARS-CoV nsp13 and SARS-CoV-2 nsp13 is more flexible and may thus confer higher enzymatic activity. To test that hypothesis, we compared the NTPase turnover rate of MERS-CoV nsp13 and SARS-CoV-2 nsp13 in an NTPase experiment with malachite green (**Fig. 6**). We performed our assays by using same enzyme concentration (i.e., 50 nM) and substrate concentration (i.e., 125 μM), and both nsp13 helicases lacked affinity tags. Similar to MERS-CoV nsp13, SARS-CoV-2 nsp13 exhibited clear preference for ATP hydrolysis over NTP and dNTP hydrolysis. As predicted, the turnover rate of MERS-CoV nsp13 was significantly slower than that of SARS-CoV-2 nsp13 for all NTP substrates, likely owing to their different hinge regions. In the particular case of dTTP, the least preferred substrate, SARS-CoV-2 nsp13 showed 11.5-fold more hydrolysis activity than MERS-CoV nsp13. In the cases of ATP and CTP, the turnover rates of SARS-CoV-2 nsp13 were greater by 3.9- and 2.3-fold, respectively. How the different NTP hydrolysis activities between MERS-CoV nsp13 and SARS-CoV-2 nsp13 correlate to nucleic acid translocation and unwinding, however, requires future in-depth analysis at the single-molecule level.

**Figure 6.**
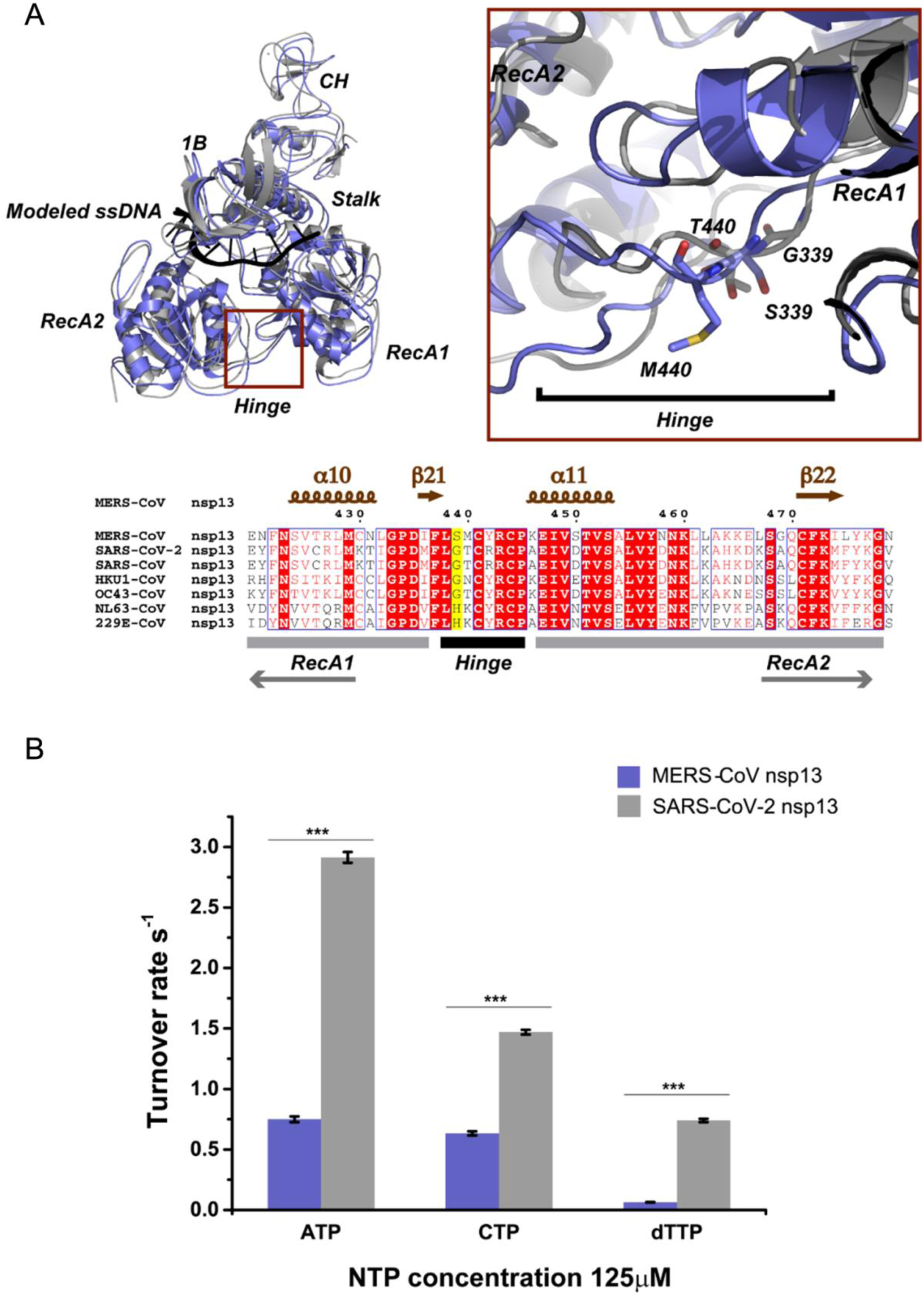
Comparison of nucleotide hydrolysis activity between different nsp13 helicases. (A) Top: Modeled structures of superimposed MERS-CoV nsp13-DNA (blue) and SARS-CoV-2 nsp13-DNA (gray); the red boxed area is the hinge region between the RecA1 and RecA2 domains of nsp13. To the right, the boxed area is magnified. Residues present major differences between MERS-CoV nsp13 and SARS-CoV-2 nsp13 are illustrated in stick models. Bottom: Structure-based multiple sequence alignment of nsp13 hinge regions of seven human CoVs. The major difference is that residue G339 of SARS-CoV-2 nsp13 is replaced by S339 in MERS-CoV nsp13, which affords the hinge in SARS-CoV-2 nsp13 greater flexibility. (B) The turnover rate (per second) of different nucleotide substrates ATP, CTP, and dTTP by MERS-CoV nsp13 (blue) and SARS-CoV-2 nsp13 (gray). For each turnover rate assessment, at least 6 independent measurements were performed to calculate the velocity of NTP hydrolysis. *** *p* < .0001.

## Materials and Methods

### Prepare of Partial Duple Oligonucleotides

Partial duplex DNA was annealed from an 18 bp single stranded DNA (Top Strand, 5’-CGA AGC TGC TAA CAT CAG-3’-Cy5) and a 37 bp single stranded DNA (Bottom Strand, Biotin-5’-TTT TTT TTT TTT TTT TTT TTT-Cy3-CTG ATG TTA GCA GCT TCG-3’), both ssDNA were purchased from Integrated DNA Technologies. Shortly, 1 μL of 100 μM Top Strand and 1 μL of 400 μM Bottom strand were mixed with 48 μL annealing buffer (containing 20 mM Tris Base, 50 mM sodium chloride (NaCl), 0.2 mM ethylenediaminetetraacetic acid (EDTA), pH 7.5) and incubated at 80 °C for 15 min and slowly cool to room temperature. The annealed DNA was diluted to desired concentrations with T50 buffer before use.

### Single Molecule FRET Measurements of DNA Unwinding by Helicase

#### Prepare of Biotinylated and Polyethylene glycol (PEG) Passivated Quartz Slides and Coverslips

The quartz slides (with drilled holes, 1 inch × 3 inch, 1 mm thick, Finkenbeiner Inc, USA) and the coverslips (24 mm × 40 mm, Corning, USA) were coated with biotin -polyethylene glycol (biotin-PEG) and PEG in order to eliminate nonspecific binding, as well as to generate biotin-NeutrAvidin bridges on the surface. The biotin-PEG and mPEG-succinimidyl valerate were covalently immobilized onto the slides surface according to the established protocol. Firstly, the slides were thoroughly cleaned with household detergent and MilliQ water, and then sonicated subsequently in the MilliQ water, acetone (Fisher Scientific, USA), 1 M potassium hydroxide (Fisher Scientific, USA) and methanol (99.8%, Fisher Scientific, USA) for 1 h each. Upon each sonication, the slides were thoroughly washed with MilliQ water. After that each quartz slide were burnt with a propone torch for 2 min, and immersed into the MilliQ water immediately. The slides were subsequently incubated in methanol containing with 1% (v/v) 3-Aminopropyltriethoxysilane (APTES, Sigma, USA) and 5% (v/v) acetone for 10 min, and then incubated for another 10 min after 1 min sonication. Upon incubation, the slides were thoroughly washed with MilliQ water and methanol and blow dry with air. After that 80 μL mixture of the 200 mg/mL methoxy-polyethylene glycol succinimidyl valerate and 36.67 mg/mL biotin-polyethylene glycol succinimidyl valerate (m-PEG-SVA, biotin-PEG-SVA, Laysan Bio Inc, USA) was dropped onto the treated quartz slides, and the cover slides was placed on the top of the quartz slides to form a chamber. After that, the chamber was incubated in a wet box in dark over night at room temperature. Upon incubation, the chamber was separate carefully and the surfaces were washed thoroughly with Milli Q water then blow dry with air. The PEGylation process was repeated. The flow chamber was assembled from the biotin-PEG coated quartz slide and a coverslip using double sided tape and Epoxy glue.

### Immobilization of DNA and Helicase onto the Substrate

The partial duplex DNA was immobilized onto the substrate by the biotin-NuetrAvidin bridge, and the helicase specifically bound to the junction of the ssDNA and dsDNA at the partial duplex DNA. Firstly, 50 µL of 0.1 mg/mL NeutrAvidin (Fisher Scientific, USA) was added into the channels and incubated for 5 min at room temperature. Upon incubation, the unbound NeutrAvidin was washed out with 200 µL T50 buffer for at least 3 times for each channel. Subsequently, 50 µL of 15 pM partial duplex DNA was added into the channel and incubated for 10 min at room temperature. Then any unbound DNA were washed out with 200 µL T50 buffer for at least 3 times. After that, 50 µL of 20 nM helicase was injected into the channel and incubated for another 10 min at room temperature. Upon incubation, 50 µL of NTPs at different concentration in oxygen scavenger solution (0.1 mg/mL glucose oxidase (Sigma, USA), 0.02 mg/mL catalase (Sigma, USA) and 0.8% (w/w) dextrose (sigma, USA), 3 mM 6 -hydroxy-2,5,7,8-tetramethylchroman-2-carboxylic acid (trolox, sigma, USA)) was injected into the channel. The sample was imaged immediately after the NTPs injection.

### Single-Molecule Imaging Through Total Internal Reflection Fluorescence Microscopy (TIRF)

The DNA unwinding events by the helicase were recorded with a homemade TIRF microscopy. The detailed structure of the microscopy was reported before. Specifically, the Cy5 labeled sample was excited by a 633 nm laser beam while the Cy3 labeled sample was excited by a 532 nm laser beam. Videos were taken with 100 ms exposure time and 600 frames were recorded. Obtained data was analyzed in real time using the custom software obtained from Dr. Taekjip Ha’s group at Johns Hopkins University.

## Acknowledgements

This work was supported by the Special Coronavirus (COVID-19) Research Pilot Grant Program from University of Cincinnati College of Medicine to J.D. and CRP-ICGEB Research Grant 2019 (Grant number: CRP/CHN19-02) to C.S..

## Author contributions

S.C. and J.D. designed and directed the study. X.H. performed single-molecule measurements and analyzed data. W.H., B.Q., Z.L., P.H., and R.Z. purified proteins and conducted bulk experiments. All authors discussed the results, and wrote the manuscript.

## Table of Contents artwork

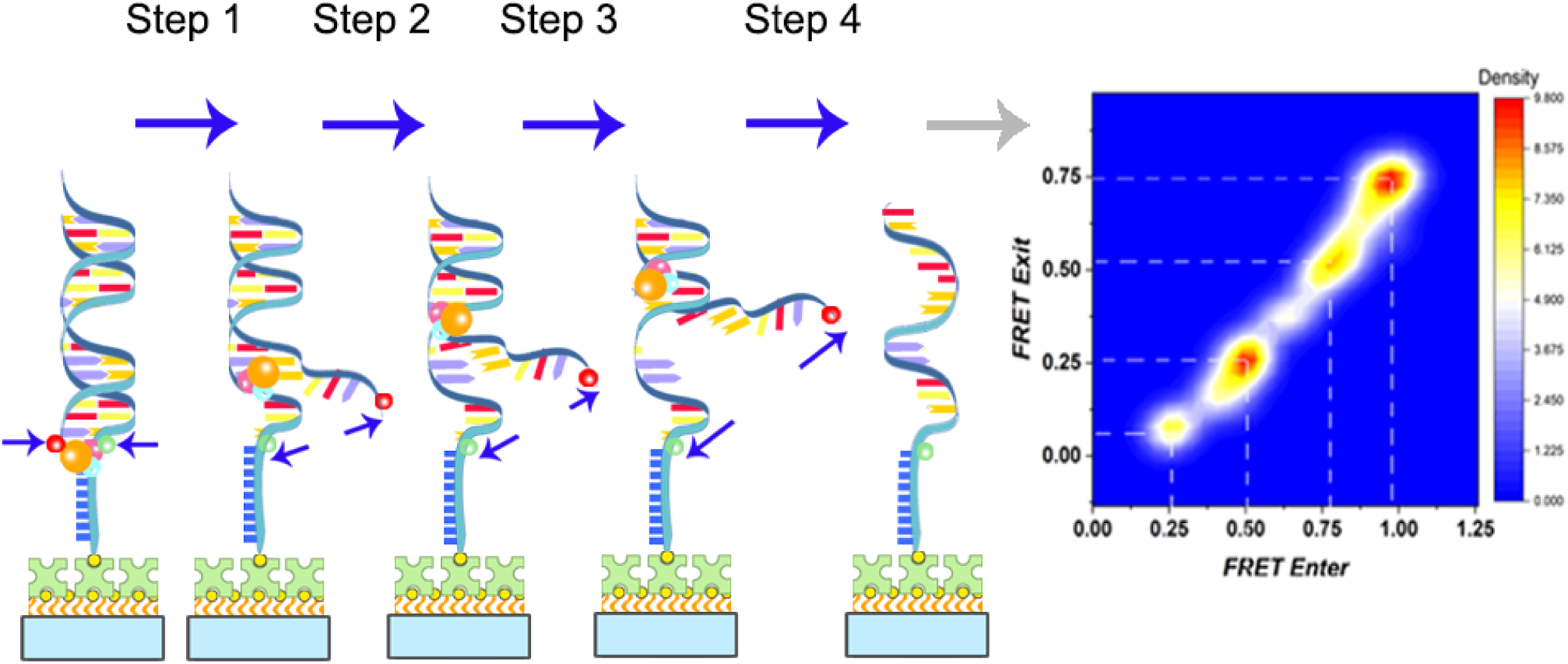

