## Supporting Figures for "Mechanism of duplex unwinding by coronavirus nsp13 helicases"

**Table S1**: The relationship between the acceptor/donor number and the ATP injection time for Figure 1.

| Time (s) | 0 | 10 | 20 | 30 | 40 | 50 | 60 |
|---|---|---|---|---|---|---|---|
| Acceptor Number | 184 | 133 | 103 | 84 | 79 | 67 | 50 |
| Donor Number | 418 | 433 | 428 | 438 | 451 | 444 | 432 |

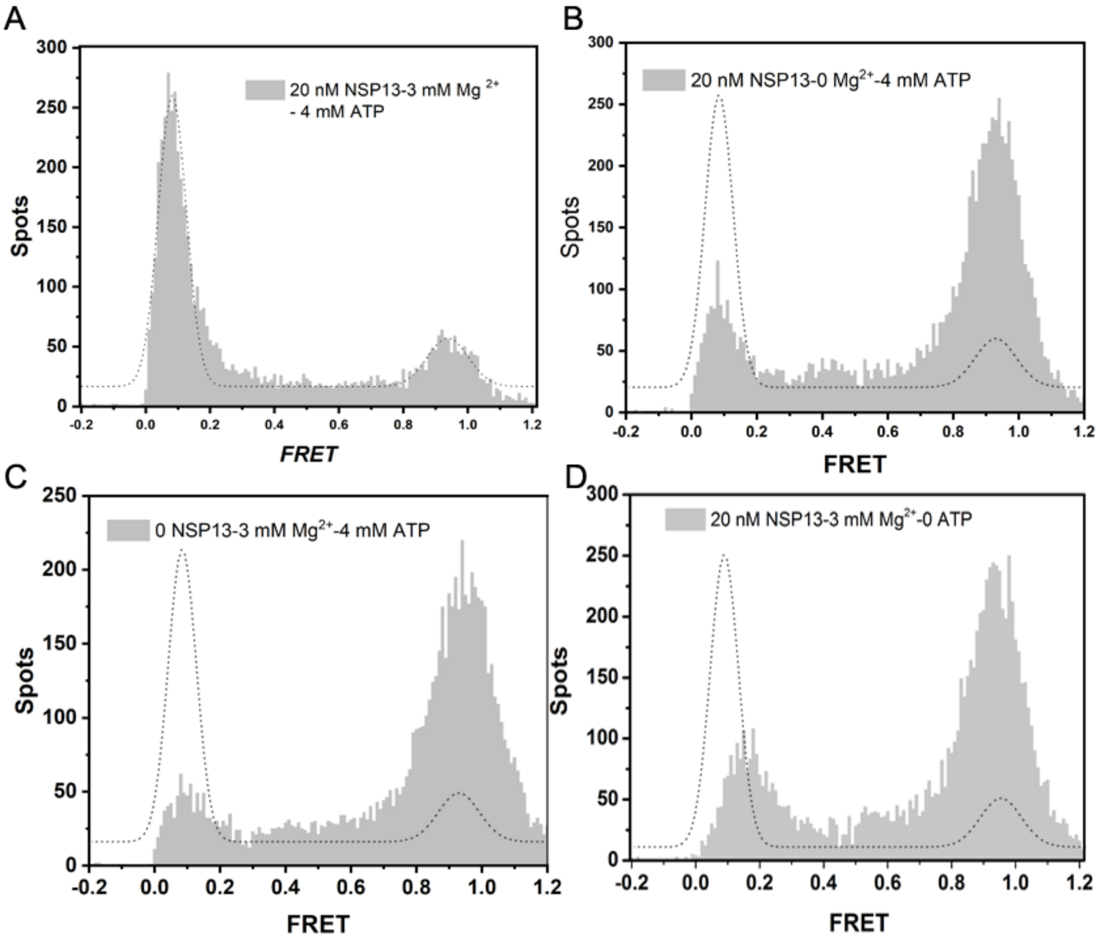


**Figure S1:** The MERS-CoV NSP13 helicase unwinding is ATP-dependent and requires all components of helicase, ATP and $Mg^{2+}$ to reach the maximum decrease in FRET values. FRET histograms of unwinding DNA (low FRET) and no unwinding DNA (high FRET) (a) by 20 nM MERS-CoV NSP13 helicase with 3 mM $MgCl_2$ after 4 mM ATP was injected for 1 min. The two peaks were fit with a Gaussian function (dotted line). (b) by 20 nM MERS-CoV NSP13 helicase with 0 mM $MgCl_2$ after 4 mM ATP was injected for 1 min. (c) 0 nM MERS-CoV NSP13 helicase with 3 mM $MgCl_2$ after 4 mM ATP was injected for 1 min. (d) 20 nM MERS-CoV NSP13 helicase with 3 mM $MgCl_2$ after 0 mM ATP was injected for 1 min.

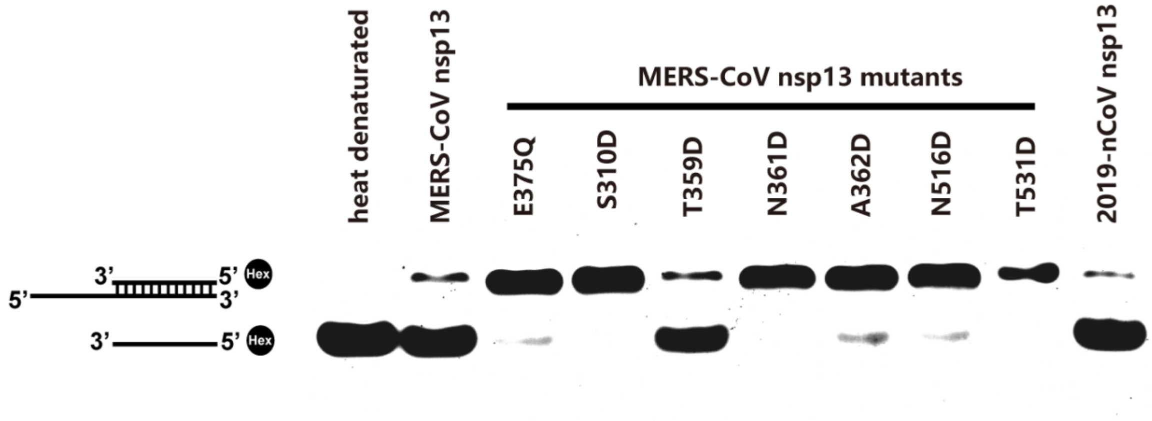


**Figure S2. Mutagenesis study of MERS-CoV nsp13 and 2019-nCoV nsp13.** Ensemble helicase assay shows that MERS-CoV nsp13 and 2019-nCoV nsp13 unwind DNA partial duplex (18-bp) with 5' overhang (10-nt). The top strand was labelled with HEX at the 5’ prime. Heat denatured controls is indicated. ATP was added as the chemical energy source. The reactions were resolved by native-PAGE and visualized by GE Typhoon Gel Imaging Scanner. Helicase activity of MERS-CoV nsp13 mutants are indicated with a line on top of the lanes. E375Q is an NTPase inactive mutant, severing the negative control. The other mutants are located across the nucleic acids binding groove formed by RecA1-A2 domains.
